# Reconstruction and dynamics of human intestinal microbiome observed *in situ*

**DOI:** 10.1101/2020.02.25.964148

**Authors:** Xiaolin Liu, Min Dai, Yue Ma, Na Zhao, Huijie Zhang, Liyuan Xiang, He Tian, Guanghou Shui, Faming Zhang, Jun Wang

## Abstract

Gut microbiome are studied primarily using fecal samples in humans and we gained vital knowledge of compositional and functional capacities of gastro-intestinal microbial communities. Yet, fecal materials limit our ability to investigate microbial dynamics in different locations along GI-tract (*in situ*), nor in finer temporal scales as they are infrequent. With a technology developed originally for fecal material transplantation, colonic transendoscopic enteral tubing, we were able to sample ileocecal microbiome twice daily, and carried out metagenomic as well as metatranscriptomic analyses. Ileocecal and fecal microbiome are similar in metagenomic profiling, yet their active genes (in metatranscriptomes) are highly distinct. Both were perturbed after laxatives and then became more similar to microbiome prior to treatment, demonstrating resilience as an innate property of gut microbiome. Ileocecal microbiome transcriptomes sampled during day and night revealed diurnal rhythmes exist in certain bacterial species and functional pathways, in particular those related to short-chain fatty acid production. Lastly, metabolomic analysis in fecal and urine samples mirrored the perturbance and recovery in gut microbiome, indicating crucial contribution of gut microbiome to many of the key metabolites involved in host health. Our study provides interesting novel insights into human gut microbiome, and demonstrates the inner resilience, diurnal rhythmes and potential consequences to the host.

## Introduction

Investigations into human gut microbiome has revealed the pivotal role of microbial communities in host health and disease^1^, however the majority of the studied microbiome are from defecated feces, which are powerful representations of gastrointestinal (GI) microbial ecosystems yet still with limitations. Fecal microbiome largely represent the last stage of transitions of microbial communities along the GI tract, and as numerous biogeographical analyses revealed the distinction of fecal microbiome from different GI sections^2^, where important biological processes take place. Largely composed of bacteria and archaea from lumen (content), fecal microbiome also indicate poorly on the mucosal microbiome, community at the intimate interface of cross-talk with the host and especially interactions with immune systems^3, 4^. In addition, continual fecal samples are rarely taken within short intervals but mostly daily due to human physiology, which disables investigations into community dynamics with finer resolution, for instance diurnal cycles that are feasible in mouse and have already been shown to be closely linked to host physiological homeostasis^5, 6, 7^.

While most of the time-series studies of human gut microbiome avoid intentional challenges or stress, perturbations in human gut microbiome are common and impact their composition and functions^8, 9^, and the shifts as well as dynamics are commonly indicative of community robustness under environmental stress. Extreme dietary changes are linked to quick responses of human gut microbiome in healthy individuals^8, 9^, while longer-term dietary habits are shown to be associated with more generalized microbiome clusters known as enterotypes^10, 11^. Gut microbiome response distinctively towards many classes of antibiotics^12, 13^, and the effects can sometimes detectable after months or years since last administration, though such studies are usually retrospective^14, 15, 16^. Continuously monitoring of human gut microbiome after infection or medical treatment has achieved interval of days after the event or interventions^17^, and study into one type of large-scale replacement of gut microbiome, fecal microbiota transplantation (FMT), demonstrated that an foreign microbiome can partially establish and replace the original gut microbiome, resulting in a mosaic of original and new gut microbiome^18^. Osmotic laxatives can remove large portions of GI microbiome and widely used for treating constipation but also before colonoscopy ^19, 20^, which is linked with long-term partially alterations of gut microbiota after a week of continuous administration in mice, with negative consequences on host intestinal and immunological homeostasis^5^.

Feces usually recover to previous physical state within days after osmotic laxatives in healthy individuals, as observed in clinical practices. Such reconstruction process provides a fascinating opportunity to examine the dynamics and robustness of healthy human gut microbiome^21, 22^, after strong perturbation in terms of quantitative change in microbiome. In addition, a technique of placing a tube through the anus into cecum or terminal ileum under endoscopy for repeated administration of FMTs or medications is called colonic transendoscopic enteral tubing (TET)^23, 24^. The TET tube also reversely allows sampling of fluid in intestine, facilitates analyses of both fecal microbiome after osmotic laxatives and also ileocecal microbiome *in situ*. In invited healthy volunteers adminisrating osmatic laxatives, the dynamics of reconstruction in fecal microbiome on daily basis, and ileocecal microbiome with a 12-hour interval to investigate fine-resolution dynamics that might contribute to host circadian rhythms were investigated, empowered by combined metagenomic, metatranscriptomic and metabolomic analyses.

## Results

Five healthy volunteers underwent osmotic laxatives treatment, colonoscopy and colonic TET in our study. Fecal samples were collected before taking laxatives as well as all days remaining TET (Figure 1). Ileocecal fluid were sampled each day at 10 am as well as 10 pm, washed off using normal saline solution and collected through TET using syringe. In total, metagenomic DNA as well as RNA were extracted from 28 fecal and 43 ileocecal samples for metagenomic and metatranscriptomic sequencing and analysis^25, 26^ on Illumina NovaSeq platform. Urine samples were also collected from each volunteer and together with fecal samples from the same sampling time, were analyzed for metabolome.

**Fig. 1.**
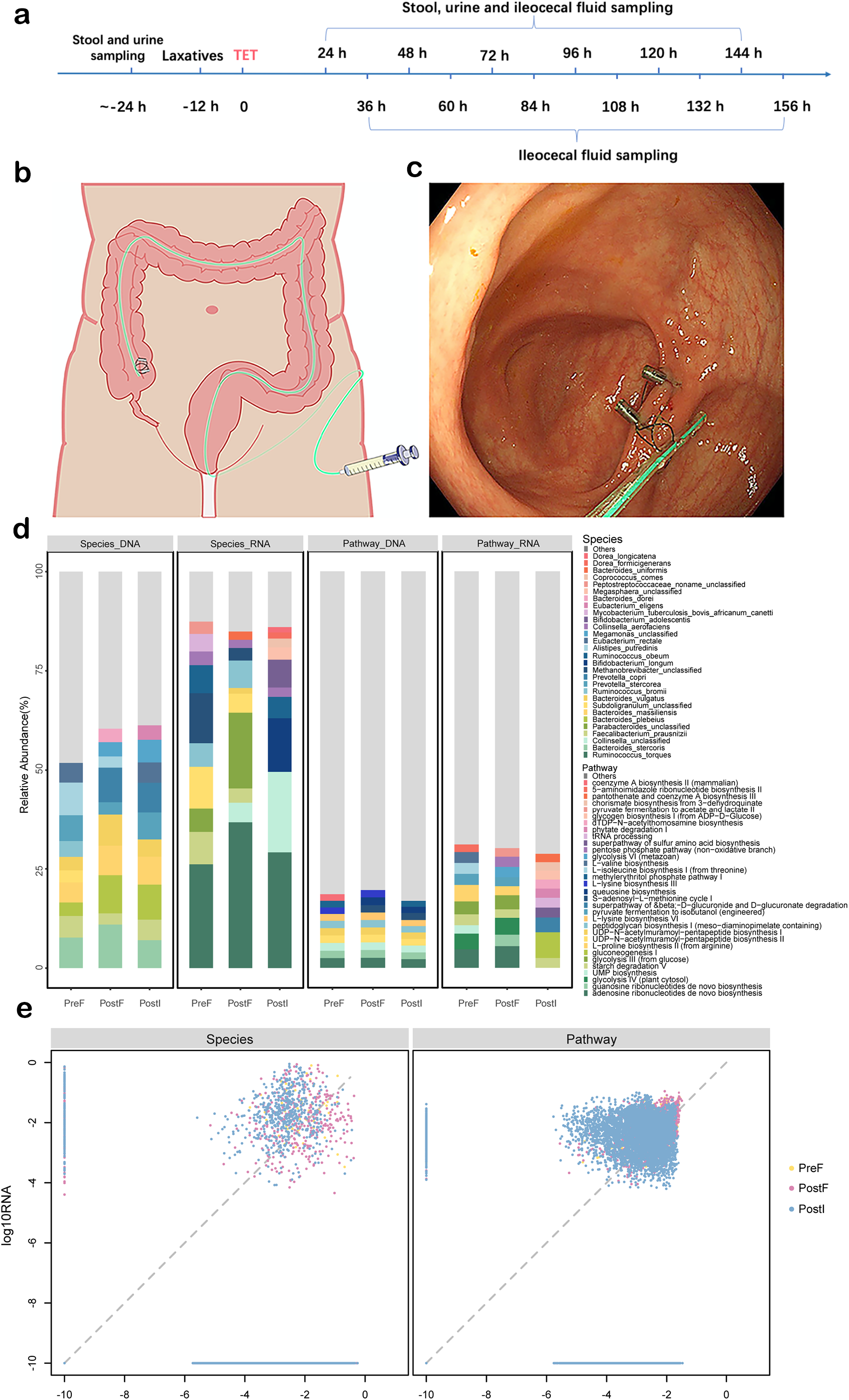
Fecal and ileocecal microbiome are distinct in composition and functionality. (a) Flow chart of sample collection. (b) Schematic diagram of colonic TET; (c) Colonoscopy view of terminal of TET tube fixed on the ileocecal valve by two endoscopic clicks; (d) An overview of relative abundance of composition at species level and functional pathways. Different colors of the bar represent different species or pathways, and the relative abundances are the average of individuals in the three group, fecal samples before osmosis (PreF), fecal samples after osmosis (PostF), and ileocecal samples after osmosis (PostI). Four panels show species composition from metagenome (Species_DNA) and metatranscriptome (Species_RNA), and functional pathways from (Pathway_DNA) and metatranscriptome (Pathway_RNA). Contributions of species and pathway were scaled to sum to 1 respectively within each group; (e) Comparison of metagenome-vs. metatranscriptome-derived species composition and functionality. X and Y show log(10) relative abundance of species and pathways in metagenome (DNA) and metatranscriptome (RNA), added a very small number(10^-10) before to aovid negative infitite values.

### Fecal and ileocecal microbiome are distinct in composition and functionality

Analysis of standing (DNA-based) and active (RNA-based) microbial communities revealed their distinct features both in defecated feces samples as well as in ileocecal samples. For fecal samples collected before osmotic laxatives, standing communities are largely dominated by species *Alistipes putredinis, Bacteroides stercoris, Prevotella stercorea, Faecalibacterium prausnitzii, Bacteroides massiliensis* among others (Figure 1, **Figure S1)**. Yet those microbial species do not always occupy the majority in RNA reads, rather, species *Ruminococcus torques, Methanobrevibacter, Subdoligranulum*, *Faecalibacterium prausnitzii, Ruminococcus obeum* are more prominent in RNA-based surveys (Figure 1, **Figure S1)**. This could be extended for functional pathway analysis, for the most abundant pathways in metagenomic DNA are adenosine ribonucleotides de novo biosynthesis, UMP biosynthesis, UDP-N-acetylmuramoyl-pentapeptide biosynthesis II (lysine-containing), UDP-N-acetylmuramoyl-pentapeptide biosynthesis I (meso-diaminopimelate containing) and peptidoglycan biosynthesis I (meso-diaminopimelate containing) (Figure 1, **Figure S2)**. While adenosine ribonucleotides de novo biosynthesis remains the highest in abundance in metatranscriptome, glycolysis IV & III, starch degradation V and gluconeogenesis I are actually among the highest in RNA reads (Figure 1, **Figure S2)**. Similar distinction could be observed in ileocecal samples as well as fecal samples collected post-colonoscopy, that DNA-based profiling of microbial composition or functionality, even though most widely used, does not represent the actually status of the microbial communities in terms of activity. Many of the abundant taxa in DNA-based profiling were found to be with low activity in terms of RNA-transcription, while many metabolic pathways found highly expressed in metatranscriptomic analysis were very low in metagenomic (DNA) reads (Figure 1). The reverse was also true for many bacterial species as well as metabolic pathways.

Ileocecal microbiome, collected *in situ*, reveal essential differences in taxonomy and functional pathways when compared to fecal samples mainly at actively transcribed genes (metatranscriptome). At DNA level, species composion and pathways are very similar between ileocecal and fecal samples from anus, without significant differences. Yet at transcriptome level, fecal and ileocecal samples showed essential differences, *Collinsella unclassified, Bifidobacterium longum, Bifidobacterium adolescentis, Ruminococcus obeum* are highly active in ileocecal microbiome while *Parabacteroides, Subdoligranulum, Faecalibacterium prausnitzii, Bacteroides uniformis, Bacteroides vulgatus* are in fecal materials (Figure 2, **Figure S3).** In terms of functional pathways, adenosine ribonucleotides de novo biosynthesis, glycolysis IV & III and guanosine ribonucleotides de novo biosynthesis are enriched in fecal microbiome RNA, suggesting high metabolic activities, while gluconeogenesis I, superpathway of beta−D−glucuronide and D−glucuronate degradation, superpathway of sulfur amino acid biosynthesis, tRNA processing and dTDP−N−acetylthomosamine biosynthesis are potentially most active in ilealcecal samples.

**Fig. 2.**
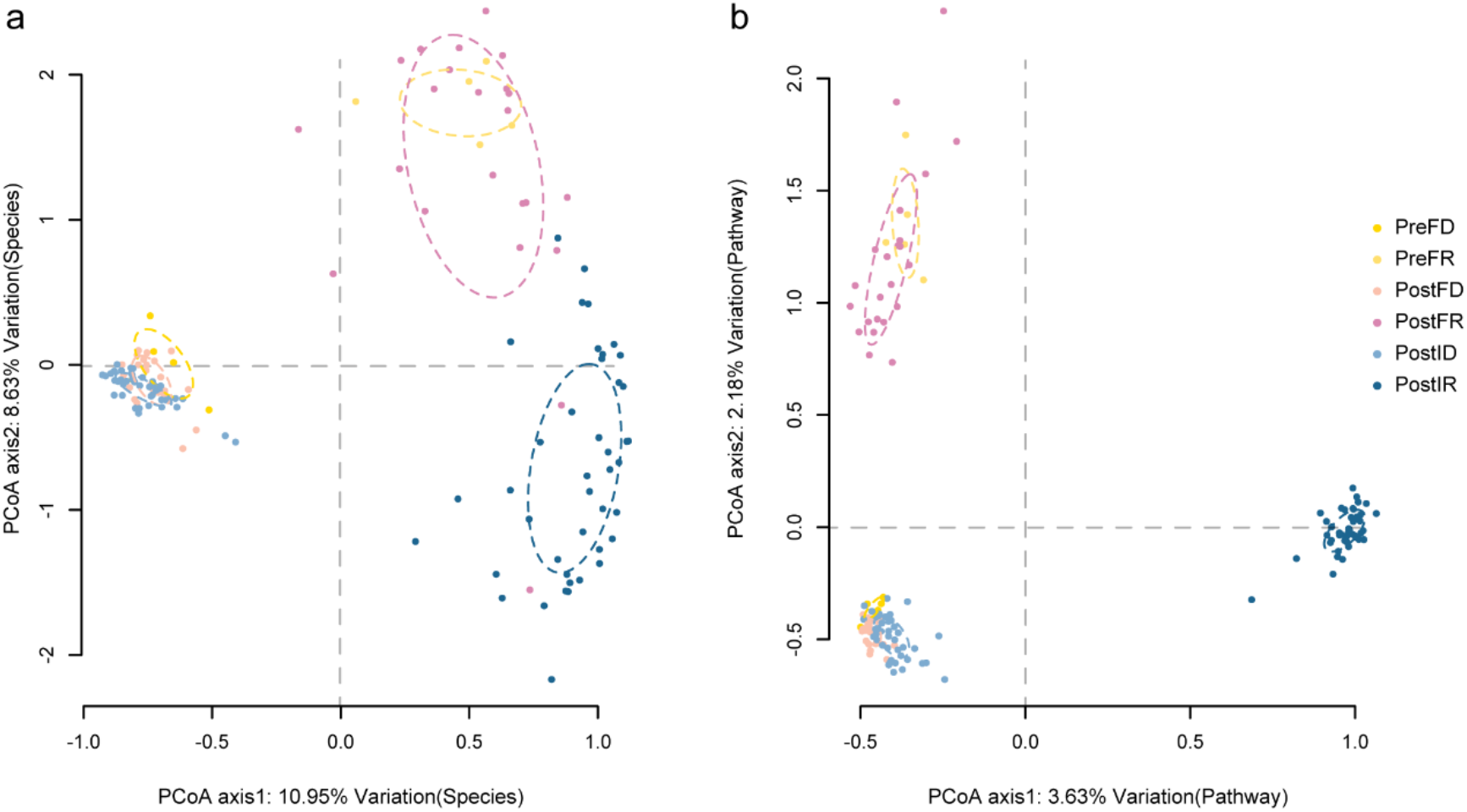
Dissimilarity between defecated feces and ileocecal samples in species composition and functional pathways. (a) The principal coordinate analysis (PCoA) of metagenomic (DNA) and metatranscriptomic (RNA) composition at species level for samples grouped by the sampling time and location (PreF, PostF, PostI), with ellipses showing 95% CI. (b) The principal coordinate analysis (PCoA) of metagenomic (DNA) and metatranscriptomic (RNA) functional pathways for samples grouped by the sampling time and location (PreF, PostF, PostI), with ellipses showing 95% CI.

### Fecal microbiome dynamically reconstruct after osmotic laxatives treatment

With strong perturbation of osmotic laxatives, fecal microbiome of each volunteer all showed dynamic recovery/reconstruction in the days following colonoscopy. Earliest fecal samples after laxatives showed differences in bacterial composition, both at DNA and RNA level. In DNA-based analysis, the most dominant taxa in the earliest fecal samples post-laxatives are *Bacteroides plebeius*, *Bacteroides massiliensis*, *Bacteroides stercoris, Bacteroides vulgatus* and *Megamonas unclassified*, instead of *Alistipes putredinis, Bacteroides stercoris, Prevotella stercorea, Faecalibacterium prausnitzii* and *Bacteroides massiliensis* in the fecal samples before **(Figure S4)**. This points to either different sensitivity to laxatives among different species in the process of being washed out, or variations in growth/recovery within the short period of laxatives to next episode of defecation. Active members, suggested by RNA analysis, also showed that the most active species in fecal samples changed from *Ruminococcus torques, Methanobrevibacter unclassified, Subdoligranulum unclassified*, *Faecalibacterium prausnitzii* and *Ruminococcus obeum* before laxatives to *Parabacteroides unclassified, Ruminococcus torques, Collinsella unclassified, Collinsella aerofaciens* and *Dorea longicatena* immediately post-laxatives **(Figure S4)**. Laxatives also caused shifts in abundances of metabolic pathways in DNA and RNA based analysis. The most abundant pathways changed to adenosine ribonucleotides de novo biosynthesis, UDP-N-acetylmuramoyl-pentapeptide biosynthesis II (lysine-containing), queuosine biosynthesis, guanosine ribonucleotides de novo biosynthesis and S-adenosyl-L-methionine cycle I in DNA and adenosine ribonucleotides de novo biosynthesis, guanosine ribonucleotides de novo biosynthesis, glycolysis IV & III and starch degradation V in RNA, instantly after laxatives (Figure 3, **Figure S4)**. In the following days, fecal metagenomics and transcriptomes showed trends of recovery in four out of five volunteers, as demonstrated by the dissimilarities of metagenomic as well as transcriptomic tend to decrease with time, and both principle coordinate analysis (PCoA) as well as compositional analysis could show that after drifting further from the pre-laxative fecal microbiome at certain time points, post-laxative fecal microbiome tend to become more similar to microbiome before osmotic laxative treatment (Figure 3, **Figure S5)**. Interactions between members of microbiome, as well as host physiological factors could both contribute to the recovery of microbiome after perturbation, and confer the individualized resilience of microbial communities.

**Fig. 3.**
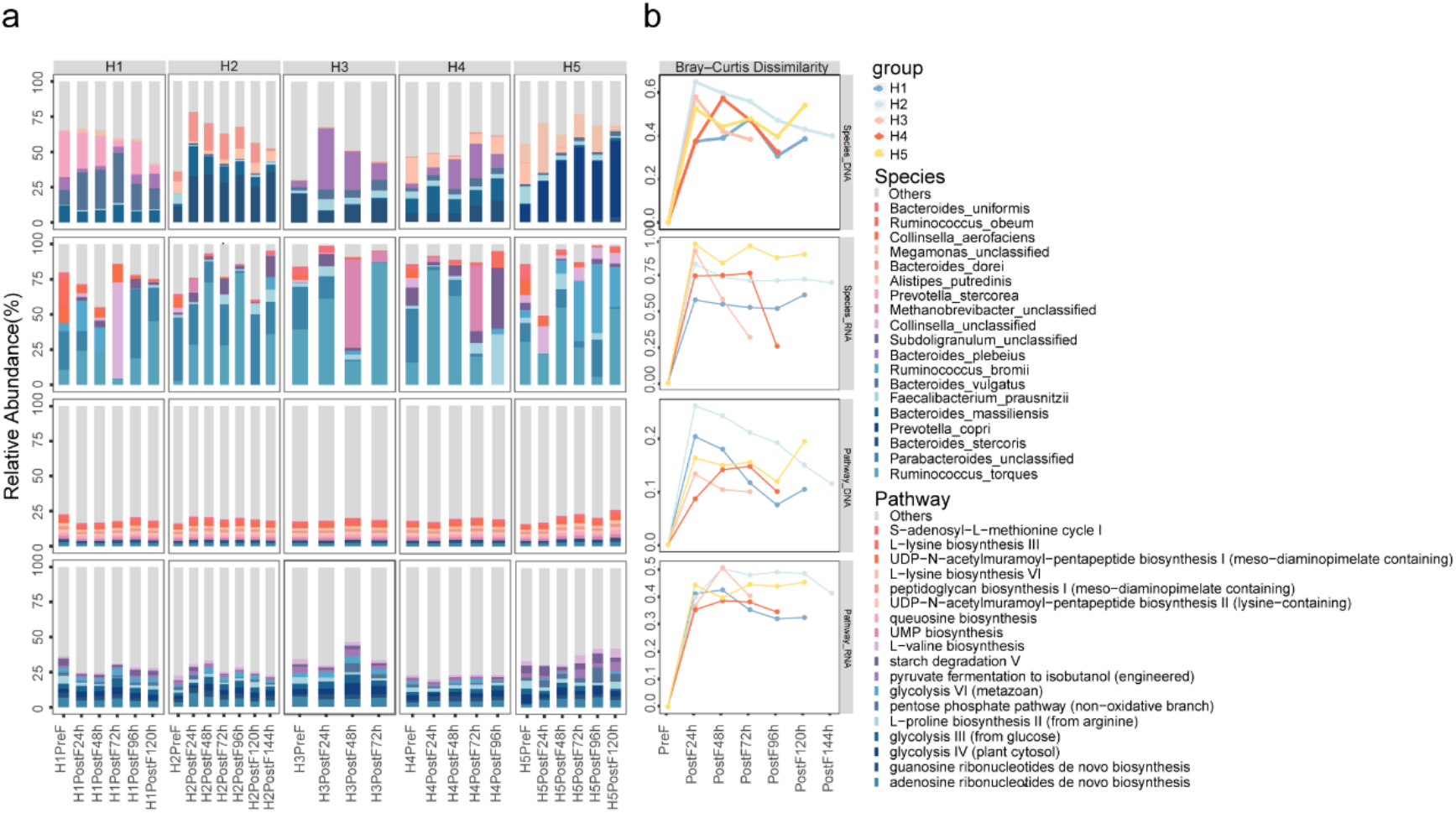
Dynamic reconstruction after osmotic laxatives of composition and function in fecal samples. (a) Bar plots show the ten most major species and pathways with the highest relative abundances in each individual (H1, H2, H3, H4, H5) at each sampling point, including before osmosis (Pre), and every day after osmosis (Post). (b) the Bray-Curtis dissimilarities from the fecal samples after osmotic laxatives (Post) to the fecal samples before interference (Pre) in each individual.

As ileocecal microbiome could not be analyzed without colon cleaning and the remained TET tube, it might be impossible to measure the recovery of microbiome using ileocecal content samples, yet their dynamics were also prominent and because of sampling intervals, could be studied with finer temporal resolution than fecal samples. Ileocecal contents, even though distinct in composition and functionality from fecal samples from the same individual and from the same day, mirror the dynamics of fecal microbiome to a large extent and could indicate that the ileocecal microbiome shifts eventually contribute to the dynamics observed in fecal samples (Figure 4). Major species determined by DNA and RNA analysis, as well as dominant metabolic pathways showed similar changes in ileocecal samples and defecated fecal samples, for instance in terms of microbial taxonomy, *Bacteroides caccae, Bacteroides uniformis, Coprobacter fastidiosus* and *Parabacteroides merdae* in metagenomic reads, *Subdoligranulum* in metatranscriptomic reads (Figure 4), and superpathway of histidine (purine) and pyrimidine biosynthesis, superpathway of pyrimidine ribonucleosides degradation, superpathway of UDP-glucose derived O antigen building blocks biosynthesis, TCA cycle I and pyruvate fermentation to propanoate I pathways in metagenomic reads, and L-isoleucine biosynthesis II, starch degradation V, glycolysis III, glutaryl-CoA degradation and purine ribonucleosides degradation pathways in metatranscriptomic reads (Figure 4). The relative dissimilarity between ileocecal and defecated feces samples before laxatives of each volunteer showed trends of decrease in three out of five individuals in at least one of the metagenomic/metatranscriptomic measures, mirroring that of the fecal samples collected post-laxatives (Figure 4, **Figure S6)**. It could be assumed that the ileocecal microbiome were also recovering to a similar taxonomical/functional composition to those before laxative treatments, thus demonstrating its own resilience towards perturbations^21^.

**Fig. 4.**
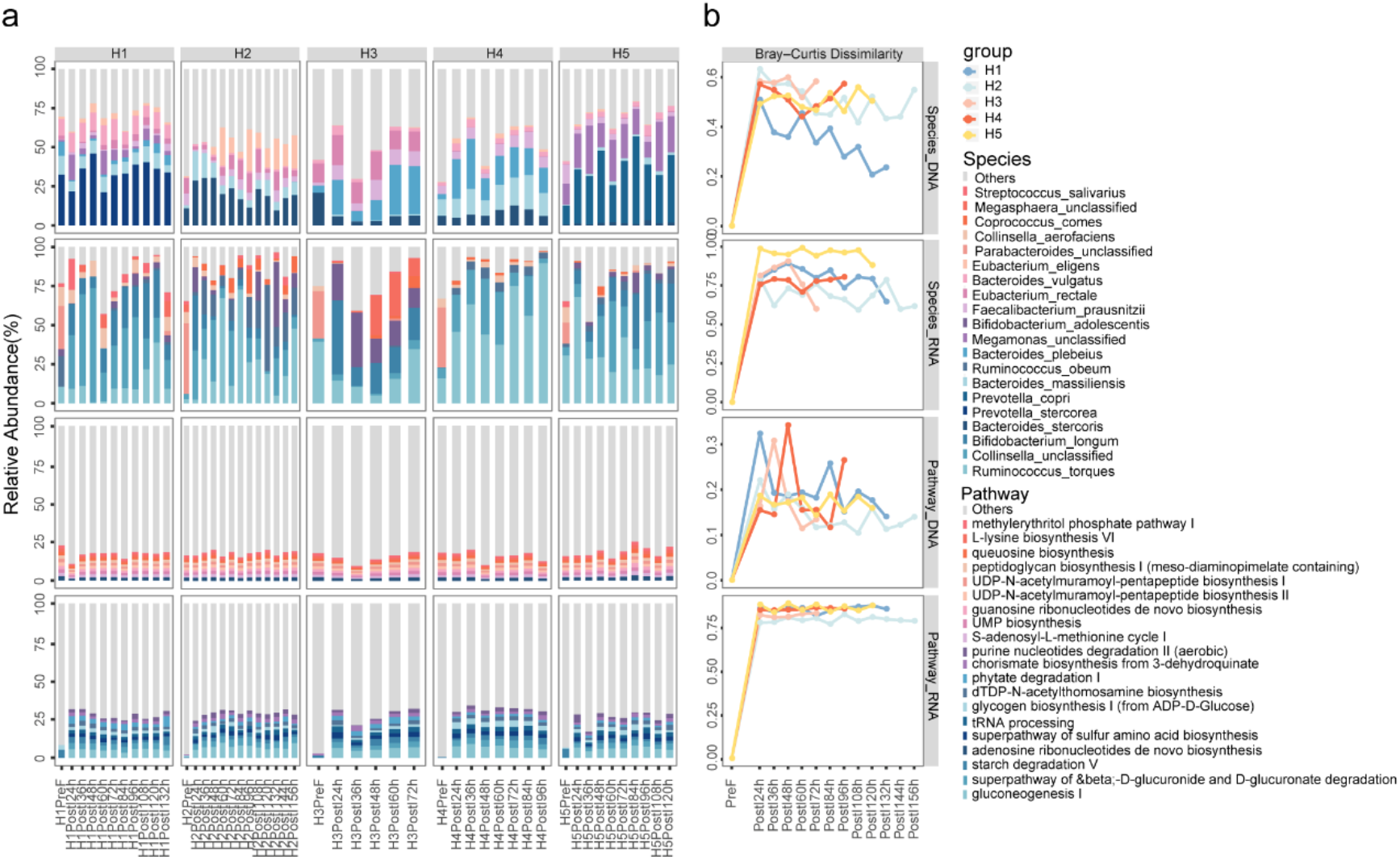
Dynamic reconstruction after osmotic laxatives of composition and function in ileocecal samples. (a) Bar plots show the major ten most major species and pathways with the highest relative abundances in each individual (H1, H2, H3, H4, H5) at each sampling time, including fecal samples before osmosis (PreF), and ileocecal samples at every 12 h after osmosis (PostI). (b) The Bray-Curtis dissimilarities from the ileocecal samples after osmotic laxatives (PostI) to the fecal samples before interference (PreF) in each individual.

### Bacterial composition and functional pathways show diurnal rhythms in cecum

The feasibility of collecting human ileocecal microbiome samples with 12-hour interval provides a unique opportunity to investigate the diurnal rhythms of GI microbiome, and more importantly *in situ.* With samples from ileocecal collected at 10 am. and 10 pm. each day through TET tube, we were able to investigate the dynamics of species composition and metabolic pathways at both DNA and RNA levels. First, based on taxonomical estimation of different bacterial species in metagenomic and metatranscriptomic analysis, it is surprising that the overall community structure has detectable differences between samples collected during the day and those collected in the evening (Figure 4, **Figure S7)**. Furthermore, we observed different species showing regular oscillations in terms of DNA abundances in each volunteer, among which a few were shared by two or more individuals: *Streptococcus parasanguinis*, *Dorea longicatena*, *Propionibacterium acnes*, *Ruminococcus lactaris, Lachnospiraceae bacterium_5_1_63FAA* and *Haemophilus haemolyticus.* While DNA-based abundance estimation reveals cell number shifts of different taxa, RNA-based analysis on the other hand demonstrates that many species have diurnal rhythm in overall gene expression (or transcription activities), namely *Coprococcus comes, Bifidobacterium pseudocatenulatum*, *Ruminococcus lactaris, Ruminococcus gnavus, Ruminococcus torques, Subdoligranulum* and *Mitsuokella*, the differences in those bacteria with diurnal rhythm among volunteers indicate a personalized trait of microbiome dynamics in the ileocecal microbiome, as well as different reproduction rate/metabolic activities of various taxonomical groups.

We further screened for pathways with prominent diurnal rhythms, primarily within transcriptomic data. This analysis again revealed individualized oscillation patterns of metabolic pathways, revealing four to 41 such pathways in each volunteer (Figure 5). Among those, purine nucleobases degradation I, coenzyme A biosynthesis II, L-lysine biosynthesis I & III, 5-aminoimidazole ribonucleotide biosynthesis I, sucrose degradation III (sucrose invertase) and superpathway of L-lysine, L-threonine and L-methionine biosynthesis II are shared by two or more volunteers, while the rest are confined to only one individual. In comparison, corresponding metabolic pathways in metagenomic data did not show a significant oscillation pattern, suggesting it is mainly the regulation of transcription in those pathways underlies the diurnal pattern, instead of the increase or decrease in genetic copies in metagenome. A general category of metabolic pathways is related to the production of short-chain fatty acids (SCFAs), including pyruvate fermentation to acetate and lactate II and acetyl-CoA fermentation to butanoate II, as well as acetylene degradation in metatranscriptome data. Those metabolic pathways are generally higher in activity at daytime, while lower at night.

**Fig. 5.**
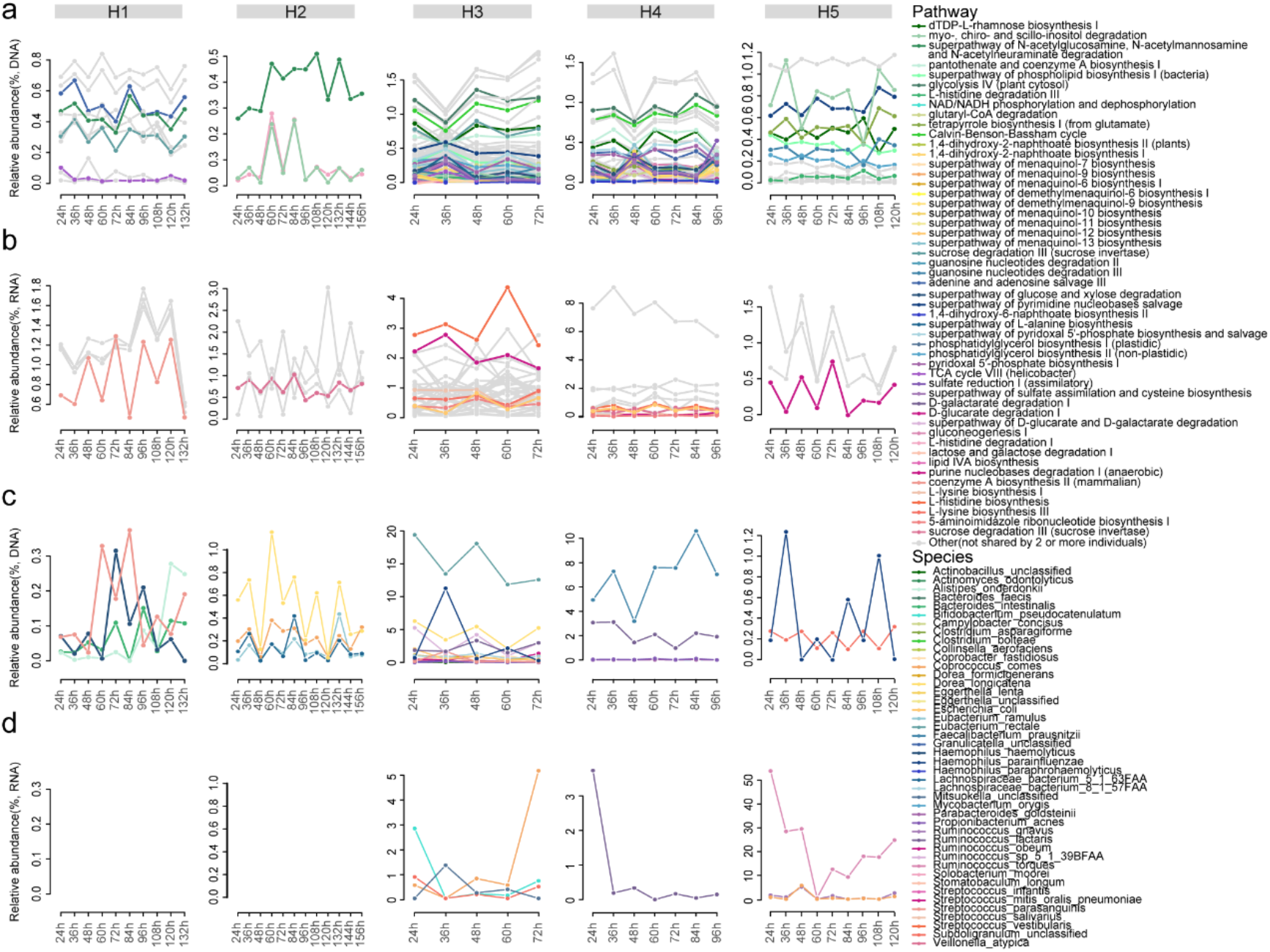
Circadian rhythm of functional pathways and bacterial species of gut microbiome in ileocecal samples. Metabolic pathways (a for metagenomic data, b for metatranscriptome data) and species (c for metagenomic data, d for metatranscriptome data) with diurnal oscillations were selected by monitoring changes in their relative abundance. Moreover, these pathways shared by two or more individuals were colored, and the pathways with diurnal oscillations shown in only one individual were displayed in grey. In contrast, all species with diurnal oscillations due to the small number of species.

### Fecal and urinary metabolome reconstruct with microbiome

To further demonstrate the potential effects of disturbance as well as recovery of gut microbiome on the host, we investigated the fecal and urinary metabolome collected before laxatives treatment and along fecal sampling after colonoscopy. Firstly, fecal metabolome demonstrated overall differences after colnoscopy (**Figure S8)**, and significant decrease in abundance due to ostomic laxatives, especially in the proportion of citrulline, N-acetyl-L-methionine, L-glutamic acid, N-acetylglutamic acid, allochenodeoxycholic acid, linoleic acid, oleic acid, 5-hydroxyindoleacetic acid, pantothenic acid, xanthine, oxypurinol, 4-trimethylammoniobutanoic acid and lithocholic acid (VIP value > 1, p < 0.05, see methods). (Figure 6). Metabolome in fecal samples also reconstruct dynamically after colonoscopy and largely correlated to microbiome recovery, resulting in a relatively similar metabolome compared to fecal samples before the disturbance **(Figure S8)**. Analyzing fecal samples post-colonoscopy revealed many associations between bacterial taxa/metabolic pathways and metabolites, however, to different degrees: we found that the composition of active species (determined by metatranscriptome data) has the strongest association with metabolome shifts, as revealed by mantel test (bray-curtis distance of species composion vs bray-curtis distance of metabolites, r = 0.38, p = 0.001), while much lower and non-significant association is found to between DNA-based bacterial composition and fecal metabolites (r = 0.13, p = 0.085). In summary, fecal metabolites are also perturbed during ostomic laxatives treatment and reconstruct together with gut microbiome afterwards, which in turn could consequently affect metabolism of the host.

**Fig. 6.**
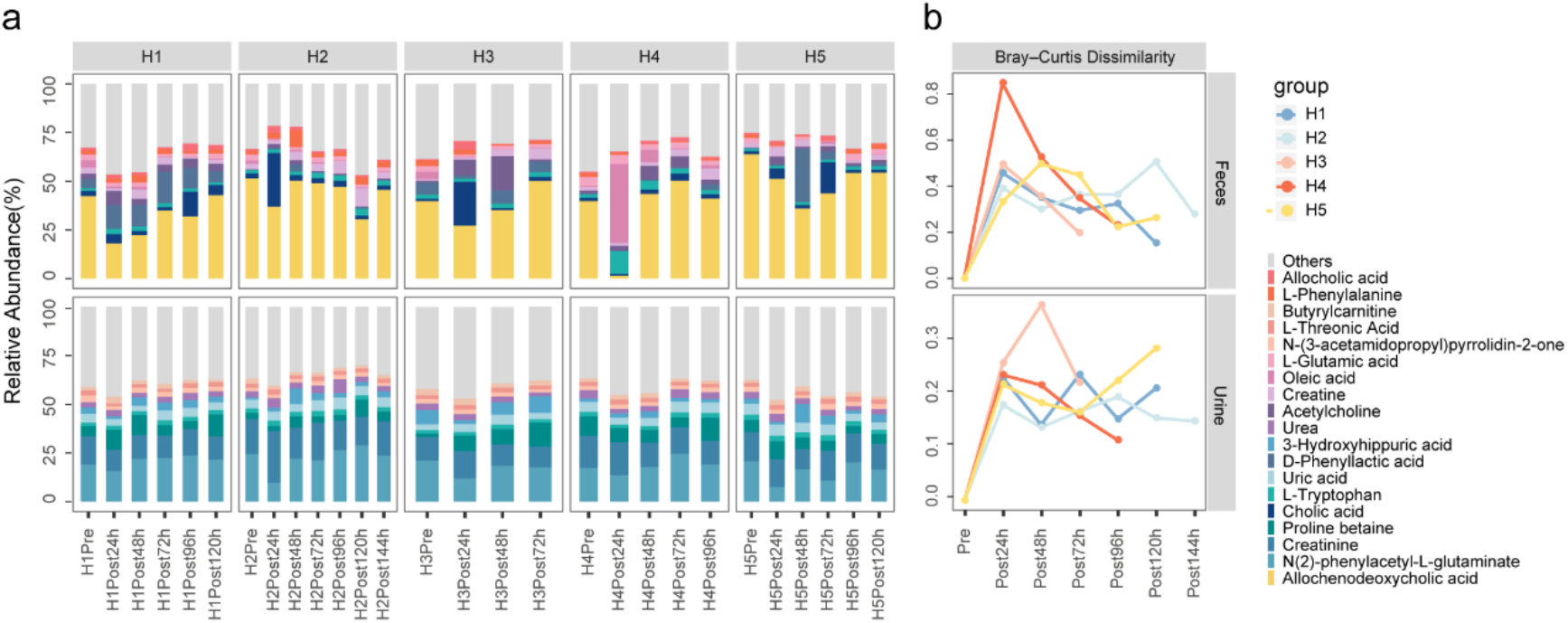
Dynamic reconstruction after osmotic laxatives of metabolites in fecal and urinary samples. (a) Bar plots show the major ten metabolites with the highest relative abundances in each individual (H1, H2, H3, H4, H5) at each sampling time, including fecal and urinary samples collected before (Pre) and after (Post) osmosis; (b) the Bray-Curtis dissimilarities from the fecal or urinary samples after osmotic laxatives (Post) to samples before interference (Pre) in each individual.

Microbiome also contribute to changes of urinary metabolome composition as well as reconstruction, although to a lesser degree than that of fecal metabolome (**Figure S8**). With depletion of gut microbiome, urinary metabolome did not have so many metabolites that have decreased, and the most promiment metabolite only has near-significant decrease (7-Methylxanthine, vip > 0, p = 0.087). In urine samples collected after colonoscopy, metabolome demonstrated lesser yet detectable recovery, mirroring the shifts in gut microbiome (**Figure S8**).Associations between metabolites and microbial taxa/metabolic pathways could also be established, with niacinamide, 1-3-dimethyluric acid and 4-trimethylammoniobutanoic acid overlapping between urinary metabolome and fecal metabolome **(Figure S9)**. Here again we observed significant correlation between active bacterial species composition and metabolome, revealed by mantel test (r = 0.28, p = 0.001), which is lower than in fecal metabolome and could be the result of less direct input from microbiome together with higher influences of host metabolic activities. Thus, urinary metabolome is also changed by gut microbiome perturbance and recovery again correlates with microbiome reconstruction.

Further enrichment analysis of the metabolites decreased in feces indicated the potential importance of microbiome-derived metabolites in many physiological and pathological conditions. Using MetaboAnalyst^27^ we discovered that changes in those metabolites are significantly associated with many metabolic syndromes including Hartnup disease, Short-Bowel disease, among others; and more intriguingly they are indicated in central neural system disorders including seizures and Schizophrenia, re-affirming the functioning of gut-brain axis is largely via various microbial metabolites (**Figure S10**). Enrichment results also include “Metabolites affected by diurnal variations” (10-fold enrichement, 15^th^ in terms of significance value), and since we established that part of those metabolites are largely dependent on the presence of gut microbiome, diurnal variations in gut microbiome can be assumed to be underlying the diurnal variations in the metabolites themselves.

## Discussion

In the present study, we collected fecal samples as well as *in situ* ileocecal samples continually after laxatives treatment and colonic TET tube that was originally designed for colonic delivery of fecal microbiota or medications. Combining metagenomic, metatranscriptomic and metabolomic analyses, we profiled the composition, function and dynamics in multiple layers of human gut microbiome, and especially revealed the individuality of reconstruction in microbiome composition and functions, which overall showed shared characteristics of internal resilience of gut microbiome. The feasibility of sampling ileocecal microbiome *in situ* and at fixed time points that are more frequent than defecated feces, provides an unique insights into the diurnal patterns or circadian rhythms in human gut microbiome for the first time. We managed to conclude that such rhythms occur at whole community level in terms of specific bacterial groups and metabolic pathways.

Bacterial species as well as metabolic pathways can have distinct patterns when they are examined at DNA (“standing”) level and RNA (“active”) levels^25, 26^. In defecated fecal samples, a collection of bacterial species including *Ruminococcus*, *Subdoligranulum* and *Faecalibacterium* are highly active in terms of transcription and potentially metabolism, and they belong to the increasingly well-appreciated group of butyrate producers, and butyrate is known to be both metabolically as well as immunologically crucial for the host, especially in the context of several metabolic as well as auto-immune disorders ^28, 29, 30, 31^. Except the top pathway being adenosine ribonucleotides de novo biosynthesis that is related to ATP production and thus energy cycling in both DNA and RNA, metabolic pathways related to nutrients metabolism (for instance glucose metabolism) are highly active in transcriptome data, even though pathways related cellular structures (for instance, components of cell walls) are most abundant in DNAs, suggesting a heterogeneous regulation of transcription in pathways aimed at different parts of microbial physiological activities.

After colon cleaning using laxatives, fecal microbiome first showed dramatic changes in composition and functionality, but in the following time points gradually reduced the distinctions to the microbiome before; considering the strong associations between cecal samples collected with fecal samples at the same time, it could be concluded that the cecal microbiome were dynamically reconstructing as well. Such construction has both the characteristics of high individuality, that different bacterial species/metabolic pathways recover to more similar abundances in each individual; and the shared characteristic of microbial resilience, that they became more and more similar in general to the microbiome pre-laxatives. Many studies have showen that the long-term stability and resilience of gut microbiome after acute disturbance including dietary intervention, medication etc^21, 32, 33, 34^; and microbiome after one-dosage of laxatives is yet another demonstration of such resilience, residing in the internal mutual interactions of microbial species^35, 36, 37^.

Regular, twice daily sampling of ceccal samples provides interesting insights into the diurnal patterns of human GI microbiome, for fecal samples are usually available on once per one-two days in humans. In metatranscriptome analysis of cecal samples, butyrate-producing bacteria again compose the majority of species that showed strong signatures of diurnal patterns, potentially a response to food intake and/or host physiological shifts in the GI tract^38^. Metabolic pathways related to production of SCFAs are among those with most prominent diurnal patterns, and those SCFAs have been recognized to be key in affecting host circadian rhythms^7, 39^. In addition, recent study has shown that circadian rhythm of host intestinal epithelial cells can be affected by MyD88-dependent HDAC3 gene expression changes, while the activities of HDAC are known to be also affected by SCFAs^40, 41, 42^. They do not only indicate replication dynamics as well as metabolic activities of bacterial groups during day or night, but might provide the eventual microbial cue for this route of gut microbiota-host cross-talk.

Contrary to the previous study in mice^5^, in which laxatives were administrated for a long-time period, volunteers in our study only received once of colon cleaning before colonoscopy and consequently the gut microbiome showed recovery instead of long-term dysbiosis; this agrees with clinical observation that colonoscopies are generally not associated with long term adverse effects in GI systems. Admittedly we did not sampling fecal samples longer enough to judge for whether gut microbiome showed total recovery, as corresponding cecal sampling is no longer feasible after TET tube falls out. The time of remaining colonic TET tube within colon generally ranged one to two weeks if using 2-4 clips for endoscopic fixaton. We are also limited on properly analyzing the ceccal sample metabolome, since the ceccal samples are wash-outs with undeterminable levels of dilutions, quantification of metabolites became difficult. And at last, host responses can be only inferred by metabolome in the fecal and urine samples, from which we did observe similar-to-microbiome recovery and re-affirmed the contribution of gut microbiome to metabolites, plus the largely microbiome-derived metabolites could be underlying important metabolic and central neural system functions, re-affirming the contribution of gut microbiome in host metabolism and gut-brain axis; and more direct measures including host intestinal epithelial gene expressions are not yet achievable in humans, only in experimental animals.

In conclusion, our study investigated the reconstruction of gut microbiome after laxative-caused depletion, and more importantly cecal microbiome with higher temporal resolution than before and *in situ*. We first time provided direct evidence of diurnal patterns of cecal microbiome in humans, at compositional and functional level. The dynamics of fecal and cecal microbiome, at multi-omics level, facilitates our understanding into gut microbial ecosystem itself, as well as how could they potentially affect host physiological including circadian rhythms.

